# Ten simple rules for managing high-throughput nucleotide sequencing data

**DOI:** 10.1101/049338

**Authors:** Rutger A. Vos

## Abstract

The challenges posed by large data volumes produced by high-throughput nucleotide sequencing technologies are well known. This document establishes ten simple rules for coping with these challenges. At the level of master data management, (1) data triage reduces data volumes; (2) some lossless data representations are much more compact than others; (3) careful management of data replication reduces wasted storage space. At the level of data analysis, (4) automated analysis pipelines obviate the need for storing work files; (5) virtualization reduces the need for data movement and bandwidth consumption; (6) tracking of data and analysis provenance will generate a paper trail to better understand how results were produced. At the level of data access and sharing, (7) careful modeling of data movement patterns reduces bandwidth consumption and haphazard copying; (8) persistent, resolvable identifiers for data reduce ambiguity caused by data movement; (9) sufficient metadata enables more effective collaboration. Finally, because of rapid developments in HTS technologies, (10) agile practices that combine loosely coupled modules operating on standards-compliant data are the best approach for avoiding lock-in. A generalized scenario is presented for data management from initial raw data generation to publication of result data.

## Introduction

Due to technological advances in instrumentation and sensor technologies, research data volumes are growing at a rate that outpaces that of the growth in computer storage and processing capacity. At the same time, society and research funders urge preservation of data in the interest of reproducibility and re-purposing of outcomes gathered with public funding, and public-private consortia require the same in order to advance their research collaborations. As these developments collide, knowledge institutes and companies are forced to formulate and implement coherent strategies to ensure that requirements and community needs are met in a way that resources can sustain. This report seeks to inform the formulation and implementation of such strategies as applied to the management of high-throughput nucleotide sequencing (HTS) data and the analysis thereof by establishing recommendations that are within reach of most knowledge institutes and their research partners, including private companies.

### DAta Management Terms and Definitions

*Data management* is the development and execution of architectures, policies, practices and procedures that properly manage the full *data lifecycle*^1^. Although the data lifecycle can be conceptualized in a variety of ways^2^, a recent report by the ICPSR [1] provides a suitable framework, reproduced in Figure 1, of the data lifecycle as it is commonly understood. The principal idea is that research data can participate in a feedback loop that promotes both re-analysis and verification as well as the development of novel research directions. Precondition for this virtuous cycle is that data management is planned and provisioned for throughout a research project. In phase 1, this includes the incorporation of a management plan in project development, possibly in collaboration with external data archiving service providers. In the case of archival of opaque blocks of data in essentially any format, services such as those provided by DANS^3^ or Dryad^4^ are available. For more granular archival of data, domain-specific archives such as those provided by the INSDC^5^ may be used. Phase 2 may include pilots and tests to establish protocols for documentation and data collection. Phase 3 proceeds with the application of practices established during project start-up. Phase 4 includes the management of *master data*, i.e. data that are common among many applications, as well as application-specific work files and backups in appropriate file structures. Phase 5 considers the *licensing terms* and *data formats* under which the data will be shared. In the final phase, the data are (re-)formatted in accordance with *community standards* for dissemination and are archived in a suitable permanent location.

**Figure 1.**
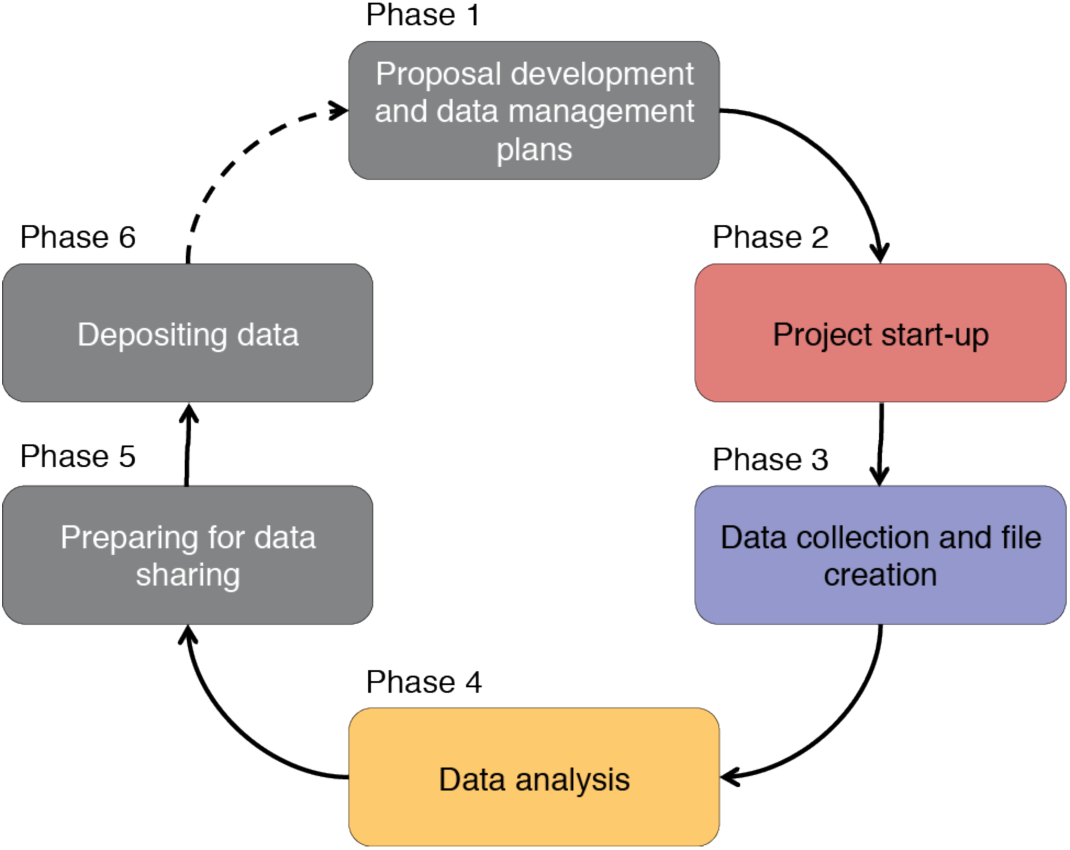
*The data lifecycle according to ICPSR*.

Within the context of research that involves high-throughput instrumentation based on sensors, data may be further classified into *raw data*, i.e. that which is directly generated by the instrument; *intermediate data*, which has undergone quality control and other data cleaning as well as normalization; and *result data*, which summarizes the results in a manner that generates knowledge. In data management, it is typically the result data that are shared in domain-specific community repositories such that they can be referred to in scholarly publications (though note the advent of alternative models in the scholarly communication cycle, such as data publications [2] and micropublications [3]), while intermediate data may be shared with project collaborators. In both cases, *metadata*, i.e. formal descriptions about data, are essential to be able to reconstruct how experiments, both the “wet” and the *in silico* steps, were performed, and to be able to steward and curate the data.

*Data stewardship* refers to the practice of storing data with provision for the required software and services to ensure its future accessibility, searching and analysis [4]. Provisioning for required software and services is needed because file formats and analytical workflows evolve rapidly, such that previously obtained results may become inaccessible or irreproducible without recourse to original software that may since have been deprecated. In this context, *open source* software may be preferred as the licensing terms of proprietary software may preclude such provisioning. *Data curation* is the related practice of ingesting, transforming, migrating, and, occasionally, disposing of data over longer timescales^6^, as informed by iterative (re-)appraisal of data, sometimes referred to as *data triage*.

### Current Trends in Biological Sequence Data Generation

In the biological sciences, a technology that strongly drives the present growth in data volumes is high-throughput nucleotide sequencing (HTS). Over the last decade, massively parallel HTS platforms have found wide adoption as relatively affordable alternatives to Sanger sequencing, albeit with their own strengths and drawbacks: the common platforms optimize different, often complementary, combinations of read quality, read length, and data volume. For example, the PacBio RSII produces very long reads suitable for scaffolding, but read quality is much lower than that of the shorter reads produced by the Illumina HiSeq, yielding a multi-stage workflow where short reads are mapped against long reads to improve precision, possibly followed by *de novo* assembly and scaffolding. Ongoing technology development will continue to disrupt workflows with the emergence of affordable, miniaturized single molecule real time sequencing such as the Oxford Nanopore MinION platform promising to yield yet longer reads more cheaply.

The relatively low and dropping cost of HTS platforms has lowered the barrier to entry for small research teams to bring sequencing to bear on a variety of research challenges ranging from “boutique” genome projects, to metagenomics, DNA barcoding, RNA sequencing, genotyping, phylogenetics and so on. By organizing into consortia, projects to re-sequence genomes at previously unimaginable scales have likewise come within broader reach. As a consequence, a recent study forecasts an ongoing shift in cost allocation away from raw sequencing towards relatively greater investment “upstream” towards experimental design and data acquisition and “downstream” towards analysis, with a relatively large but dropping expenditure at present towards data reduction and data management (Figure 2).

**Figure 2.**
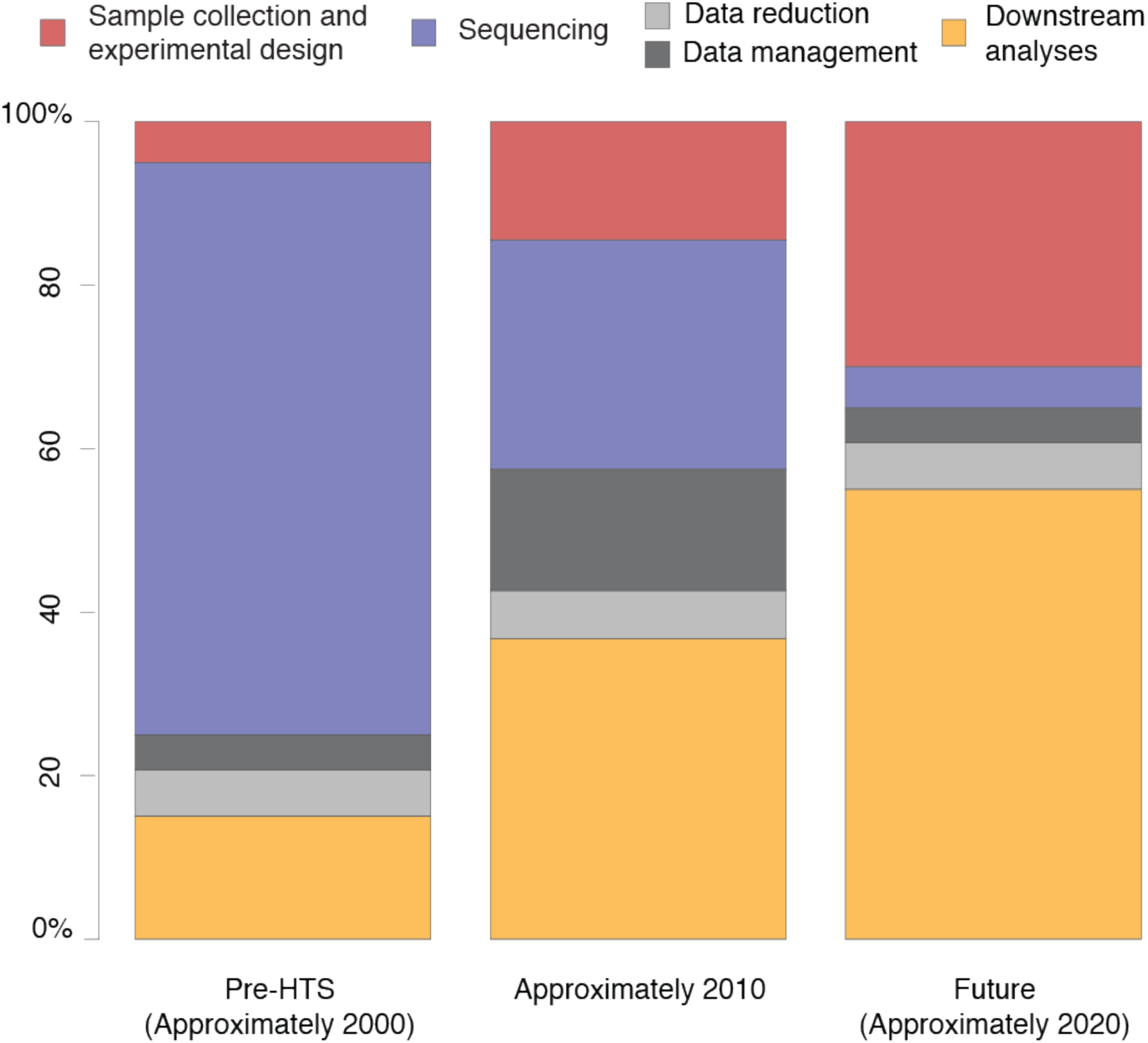
*Cost allocation outlook for nucleotide sequencing-oriented research projects [5]. The cost of sequencing itself is rapidly dropping and projected to continue to do so. Data management may be a bottleneck at present but projected to become less so in relative terms in comparison with the cost of upstream processes such as sample collection and experimental design, and downstream analysis*.

The relatively high current cost of data management in HTS projects stems not only from the large data volumes per se but also from the more complex workflow in dealing with data generated on massively parallel sequencing platforms as compared to previous technologies. Parallel data capture on current HTS platforms yields raw results - images in pyrosequencing, other types of intensities on other platforms - from which the vendor platform’s algorithms call individual bases and homopolymers at varying confidence levels. This procedure results in very many short sequence fragments that are logically organized around the instrument’s architecture and sequencing kits (e.g. according to lanes or flow cells, adaptor sequences), not around experimental design. As multiple assays are often combined in a single “multiplexed” run, an early step in data management is to sort the sequence fragments by assay. Subsequently, the sequence fragments often need to be assembled into intermediate results either by mapping against reference data or by collating overlapping fragments into larger contiguous stretches, i.e. *de novo* assembly. These intermediate results are then enriched by comparison with other, well-characterized data and by statistical analysis including variation discovery and feature prediction. At each step of this workflow, data volumes are reduced by orders of magnitude, which, if storage capacity per se forms an impediment, raises questions about the extent to which raw and intermediate data must be retained, and for how long (Figure 3).

**Figure 3.**
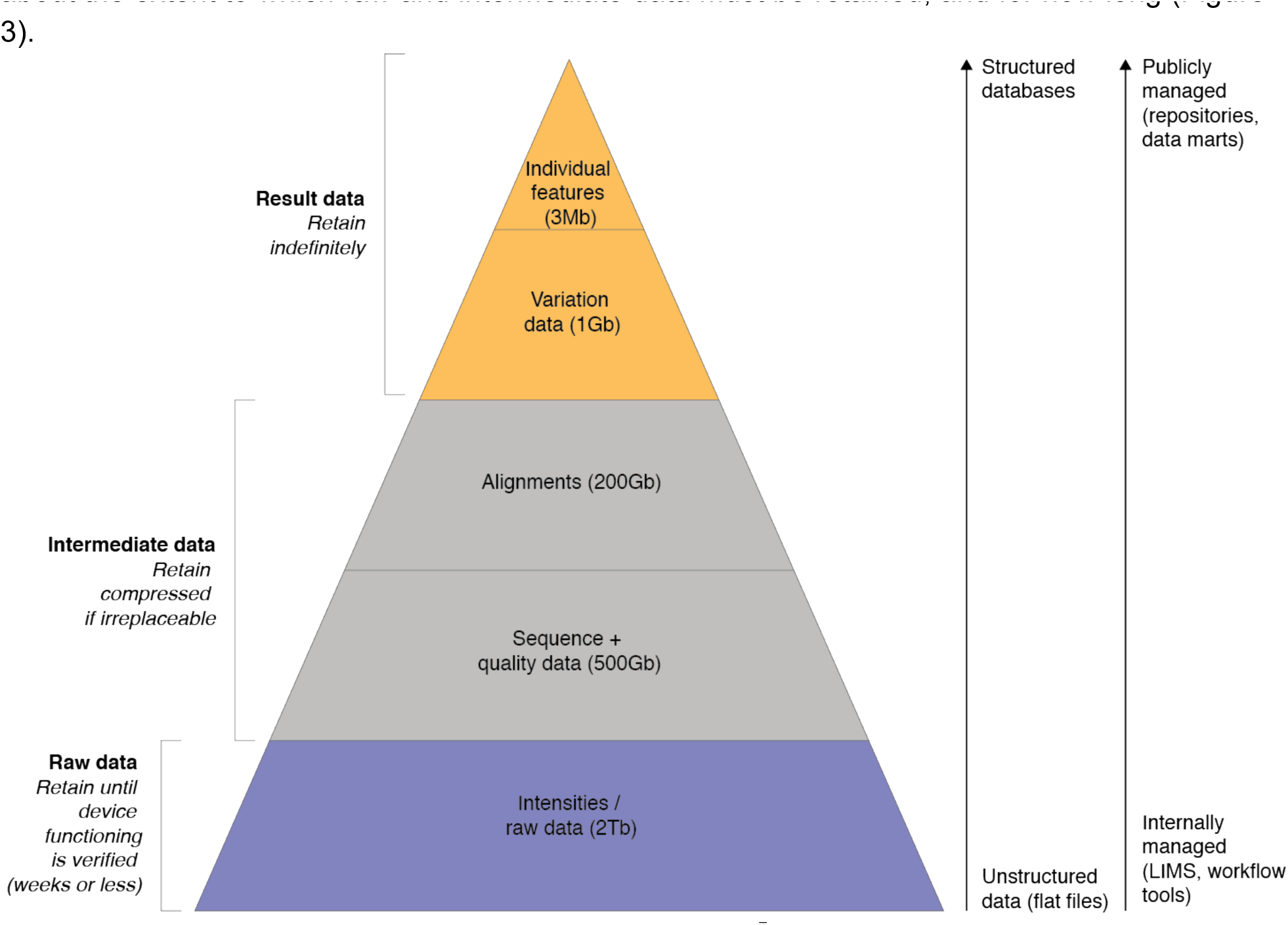
*Typical data reduction and management in HTS sequencing*^7^.

### The Accidental Sequencing Center

Due to the advent of affordable HTS platforms, knowledge institutes and company R&D departments that heretofore outsourced DNA sequencing are now finding themselves proud owners of “benchtop” (and larger) sequencing platforms, in the process having to manage complex, voluminous data sets for a large part of the data lifecycle, and potentially having to publish these to research collaborators and the research community at large.

Many knowledge institutes have found themselves to a greater or lesser extent in this position of “accidental sequencing center”. In contrast with core sequencing facilities, these institutes may on the one hand have less experience with managing large data streams, while on the other hand they are usually much more intimately involved in downstream analysis for longer stretches of time (and financial outlay). In addition, these institutes depend to a large extent on a mixture of both public funding and public/private partnerships, which each have their own requirements for data management and sharing. As a consequence, the challenges are quite formidable, and institutional policies and guidelines to cope with them are strongly called for.

Naturalis Biodiversity Center (Leiden, the Netherlands), best known as the dutch natural history museum, is one such accidental sequencing center - but many other knowledge institutes, university research departments, medical centers, and companies active in the red, green and white life sciences find themselves in a similar situation. Naturalis is a chiefly publicly-funded institute with a tangential interest in HTS as applied to a broad range of research questions, such as the genomics of non-model organisms from a broad taxonomic spectrum (including collection specimens, i.e. “museomics”), taxonomic identification through (meta-)barcoding, phylogenetic systematics, transcriptomics, and so on. Naturalis participates in a variety of research networks involving both public and private collaborators.

## Ten simple rules

This document attempts to establish ten rules in handling HTS data streams at three levels of the problem. The first level is that of *master data* management, i.e. data as they are received from sequencing, which are pre-processed, converted to an interoperable format and stored in a library location where it can be backed up. The second level is that of data analysis, where careful design and deployment of analysis pipelines and usage patterns of computational infrastructure can reduce haphazard proliferation of intermediate files while building up a “paper trail” of both the data and the analysis steps. The third level concerns collaboration among disparate research partners, where transmission of large amounts of data over the internet may be prohibitive in terms of bandwidth, and where identifiability, locatability and exhaustive documentation of metadata are key in developing effective collaboration.

The intended readership of this document is rather broad, comprising technical staff, chiefly bioinformaticians, computationally skilled researchers, ICT staff, and to a lesser extent lab staff; institutional policy makers such as R&D and ICT managers; but also cross-domain computational service providers seeking to support the computational requirements of HTS. As such, the rules are general guidelines that can be made operational in a variety of ways to fit the workflows of a variety of “accidental sequencing centers”. References in square brackets refer to endnotes (scientific literature and technical reports), superscript numbers are footnotes (links to useful information and tools on the web, clickable in digital form).

### Master Data

Master data are the core data that are consumed by a variety of applications. In the context of HTS these are sequencing reads. Because of their potential volume, these need to be managed carefully and reduced where possible. The heterogeneity of usage patterns and the potentially very large number of different applications that consume master data suggest that they must be represented in compact, interoperable form.

#### 1. Exercise data triage

At a low level, HTS platforms produce raw data files that are specific to the underlying technology. For example, pyrosequencing platforms produce digital images, while pH sensor platforms produce raw voltage data. Signal processing algorithms process these files to call nucleotide bases at varying quality levels and write the results in one of a variety of (nominally) platform-independent, community standards-compliant file formats. **Typical data triage at this level, if not already implemented in the instrument, may include purging of the underlying platform-specific raw intensities data once correct functioning of the instrument during the run has been verified**. Rough time windows cited for this include within “weeks or less”^8^ or “one month”^9^. In many cases, the next step then is to “de-multiplex” the data, for example by sorting the reads on adaptor barcode and/or primer.

The sequence and quality data that then remain contain a lot of noise whose trimming and purging may appreciably reduce data volume. **For example, short HTS reads often include stretches of very low base calling quality at the trailing end of the read, which may be truncated and discarded**. Likewise, depending on the research context, duplicate reads may be considered redundant and therefore removed or clustered. Once assembled, the sequence data may reveal yet other types of “noise”, for example as a result of contamination of the sample. Although some types of sequence contamination-such as from pathogens - might be of interest, other contaminants such as human DNA might be considered safe to discard. Careful consideration of these issues in the formulation of data triage policies (that are perhaps enforced automatically) may reduce data volume and noise considerably.

#### 2. Optimize data representation

HTS platform vendors are increasingly consolidating around well-described data standards to represent sequencing reads and their qualities, as well as alignments of sequencing reads against reference genomes. One of the common denominators is the FASTQ format [6], which is a simple flat text format that combines bases and qualities in a record-oriented layout. FASTQ may be considered convenient as, due to its simplicity, data processing scripts can quickly be developed. However, it has several drawbacks.

Firstly, plain text is relatively verbose, although a cottage industry of compression algorithms has sprung up around FASTQ to remedy this [7–9]. Secondly, FASTQ files can typically only be read sequentially, not by random access, which means that locating specific reads (e.g. mate pairs) requires scanning through files. Thirdly, a number of different FASTQ “dialects” have emerged that differ in crucial ways, including the encoding scheme for quality scores, repetition (or not) of the read identifier, and whether base and quality sequences may span multiple lines [6]. For these reasons, the binary alignment/map (BAM) format [10] may be preferred. **BAM files are a binary representation of reads and qualities, optionally mapped against a reference. Binary representations are typically more compact, and this is also true for BAM, for example in comparison with FASTQ or with the textual representation of the same BAM data as SAM, which is about four times as voluminous as BAM, on average**. Indexed, sorted BAM files can be accessed randomly like a database, which can have great advantages when processing. Also, fewer data syntax dialects exist, as many tools use the same underlying application programming interface - usually either the samtools C API, [10] or the HTSJDK Java API^10^ -, though there may be some variation in optional headers. Finally, the BAM format is one of the “container file” formats preferred by the short read archiving services provided by INSDC collaborators^11^.

A downside of BAM and other binary formats, such as the Standard Flowgram Format, SFF^12^, or the reference-based compression format CRAM [11] (which paves the way for “compressive genomics”[12]), is that their structure is more difficult to infer than flat text and that corruption such as due to truncation during network transfers may be more difficult to detect. To manage these risks, careful data stewardship (preserving the version of the API that was used) and good quality metadata, including file format descriptions and checksums, are necessary.

#### 3. Manage data replication

Unmanaged, haphazard copying of data across user home directories on workstations and compute nodes must be avoided for two reasons. Firstly, data unnecessarily strewn about multiple locations consumes space, which may be at a premium on compute nodes with expensive, fast read I/O (e.g. on solid state drives). Secondly, such proliferation creates a complex provenance trail that may be hard to reconstruct down the road. On the other hand, storage hardware will unpredictably fail, and files can become corrupted for a variety of reasons (e.g. incomplete file transfers, user errors, computer viruses), which does necessitate *managed* replication, including backups. **The simplest approach is to ingest master data into centralized storage that is backed up and from where temporary, ephemeral copies can be made to HPC resources and workstations as needed** (but see recommendation 4, below).

However, more scalable (but more complex) solutions that incorporate abstraction layers to manage metadata and access rights are gaining adoption. For example, the iRODS^13^ system is currently being used by iPlant [13], the Swedish national HTS infrastructure UPPNEX [14], the Broad Institute, the Wellcome Trust Sanger Institute, the U.S. National Center for Microscopy and Imaging Research, and the Genome Biology Unit at the University of Helsinki (for an illuminating report on iRODS deployment at these institutions, see [15]). The key appeal of this system lies in the integration of data management with arbitrary metadata, though this can be approximated to some extent by specialized analysis and data management platforms that provide more facilities for queryable annotations than the file system can (e.g. Galaxy^14^ data libraries), some LIMS systems and generic object store solutions (e.g. Amazon S3^15^ or Ceph^16^).

### Analysis

Raw sequencing data need to be analyzed in many steps in order to gain new insights. In the first instance, a number of these steps are essentially data cleaning and data organization, which produces intermediate data. Among these steps are basic quality control, e.g. to trim bases of poor quality and to filter out contamination, and assembly, either to a reference genome or *de novo*. In a dwindling number of cases this may be “enough”, but usually a large number of additional steps are performed, among which might be genome annotation, variant calling, comparisons with other samples and sequences, and a large number of statistical analyses, to arrive at the final result data. Without a planned strategy for managing these analyses a proliferation of intermediate data with unclear provenance will occur. Sensible strategies exist to manage reproducible research, to minimize movement of large data sets, and to keep a “paper trail” of the provenance of both data and analysis.

#### 4. Make analysis reproducible

During the course of a research project the same analysis steps will be executed multiple times, for example to explore a parameter space, to iterate over a set of samples, or to incorporate newly acquired data: *“everything you do, you will probably have to do over again”* [16]. This raises a number of challenges when this is done “by hand”. Firstly, people make mistakes and so without automation it is hard to ensure that exactly the same steps are followed for each iteration. Secondly, no automatic record exists of what was done. Thirdly, every time an analysis step is performed, new intermediate and result data are generated, thereby contributing to the explosion in data volumes.

To address these challenges and to make research more reproducible, broadly applicable guidelines for organizing and automating analyses have been formulated by a number of data scientists (e.g. see [16,17]). **Apart from the virtues of reproducible science *per se*, automation of analyses also aids in coping with large amounts of intermediate and result data: if the analysis can be re-run, not all intermediate steps need to be retained**. To make this possible, all the components that might influence the results of an automated analysis down to the operating system and its libraries, where relevant, should be fully specified and provisioned. In cloud computing, supporting tools to manage^17^ and provision^18^ virtual machines have become available. Moreover, recent advances in “container virtualization”^19^ and the virtualization of programming environments^20^ have resulted in a more lightweight approach to virtualization that delegates more functionality to the host operating system, thereby lowering the demands of virtualization in terms of computing power and disk space.

One level up from the operating system, analysis pipelines can be made reproducible either using specialized workflow environments^21^, declarative workflow scripting languages^22^, domain-specific languages^23^ or general purpose procedural languages^24^. Using these tools, analysis steps should be made easily (re-)runnable so that derived data can be re-generated from source.

#### 5. Bring analysis to the data

A typical workflow in bioinformatics might start by downloading data from a server to perform calculations locally on a workstation. Due to the growth in data volumes this approach does not scale well, and it is at odds with the recommendation against haphazard copying of data (recommendation 3). **Instead, the analysis workflow might be inverted: rather than bringing data to the analysis, bring the analysis to the data** [18].

Up till recently, this may have meant the provisioning and configuration of a one-size-fits-all analysis cluster or server with co-located data storage. Due to the rapid evolution of analysis tools, library dependencies and operating systems, maintenance of such systems is complex. However, **recent advances in “cloud computing” have made it possible to launch virtual machines, complete with installed analysis pipelines, on commodity hardware with co-located data storage** [19]. Analysis pipelines, even as entire virtual machines bundled with operating systems, are often much smaller in size than many types of HTS data sets, and can be tailored to a specific data set or analysis without having to modify a shared cluster or server. Even more compact ways of packaging pipelines (e.g. in “containers”^25^ or using language-specific packaging systems^26^) are available that reduce the size of analysis pipelines further.

#### 6. Track data and analysis provenance

In the course of a research project, data and analysis (e.g. custom-written scripts, workflows, configurations) undergo iterative changes. **In order to make this process fully reproducible as well as understandable this evolution needs to be recorded. In the case of small text files such as source code, summarized result data, metadata, and configuration files, version control systems such as git^27^ and subversion^28^ are strongly recommended** (e.g. see [16]), for example in combination with free cloud-based hosts^29^ that provide remote storage of the project repository.

Version control systems store incremental snapshots of the line-by-line differences between files in a repository from one revision to the next, with each revision accompanied by metadata about when, by whom, and why changes were made, i.e. a good record of the *provenance* of these files. However, because of this granularity, storage capacity can quickly be swamped especially in the case of large binary data such as BAM files, for which line-by-line differences can not be computed, which typically means that instead the entire file would be stored at each change. This suggests that the provenance of such files should be tracked separately. To keep this manageable, **raw master data files should be considered immutable, while reproduction of intermediate and result data should be automated (as per the previous section)**. This means that relatively few data versions need to be stored. Visual workflow environments such as galaxy^30^, which build up an annotatable, re-runnable history of an entire analysis^31^, may be all that is needed. Alternatively, command-line driven data management tools may be used. For example, git-annex^32^ for data integrates well with git-based analysis provenance management and can use a variety of scalable storage “remotes”^33^, while the previously-discussed iRODS system provides the flexibility to define structured metadata terms with which to record data provenance.

### Access and Sharing

Data must be accessible to internal collaborators, and (eventually) to the outside world, whether to privileged outsiders (e.g. consortium partners) or the community at large. This poses several challenges. Firstly, HTS intermediate data file sizes can be on the order of tens of gigabytes, which can be problematic if bandwidth is limited or unreliable. Secondly, once data start moving around, context may be lost and references to the data (e.g. different versions) become ambiguous. Thirdly, access may have to be controlled, e.g. for reasons of privacy or competitive advantage.

#### 7. Model data movement

Data inevitably moves between locations, even within a research institute. Without a coherent strategy for managing this movement, problems can occur. For example, sequencing instruments typically have their own limited disk space separate from institution-wide data management. If this is used for permanent raw data storage this space quickly fills up. If sequencing data are generated elsewhere, these typically arrive on physical storage media such as USB drives, which may be inaccessible to collaborators (i.e. sitting on a shelf) and may become corrupted. If the only copy of a data set is stored on a compute node, expensive storage intended as “scratch space” becomes unavailable. To prevent such problems, **data movement should be modeled and ensuing practices should be adhered to (ideally through automation)**.

A typical model might consist of the following steps: i) data are generated on an instrument or arrive on a physical medium; ii) data are ingested into cheap, networked storage; iii) checksums are computed, stored with the data, and compared with the source to ensure ingestion succeeded; iv) data and analysis are staged (see recommendation 5); v) result data are ingested and shared.

Automated management of data movement can have big benefits. For example, by implementing automated data movement at the Sanger institute, a 50% reduction in disk utilisation (out of 2PB) was achieved^34^. The movement pattern that was implemented is reproduced as the gray shaded area in figure 4.

**Figure 4.**
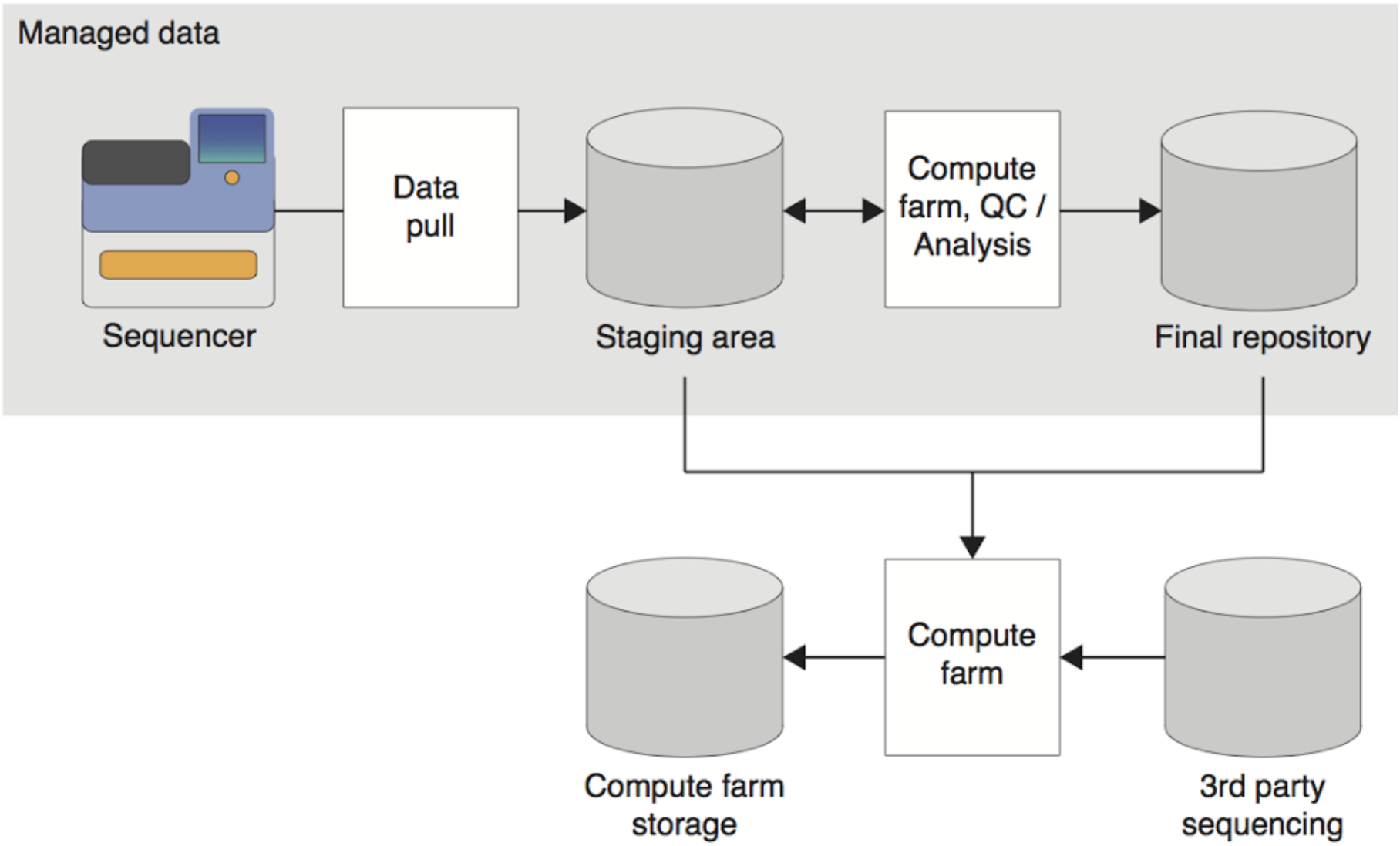
*Typical data movement pattern^35^. Raw data are automatically pulled from the sequencing platform to a staging area where quality control and analysis is performed before the data are deposited in the local, “final” repository. Data from the staging area and the repository can additionally be ingested into remote compute farms where the data are combined with that of others to perform additional analysis*.

#### 8. Make data persistently identifiable and locatable

*“The”* master data cannot solely be a file in a folder in someone’s home directory, or an email attachment sent “a few weeks ago”, or even *“the* dropbox folder” (which one?). **In collaborating and communicating with others, data needs to be uniquely, persistently identifiable and locatable**, or collaborators will have problems identifying and accessing the relevant data. For most purposes, URLs that can be referenced by metadata such as timestamps and checksums may be enough during the course of a research project, as URLs are both globally unique identifiers by virtue of the domain name system as well as locators.

Alternatively, data hosting and sharing technologies that can be queried and traversed, such as FTP servers or object stores, can be used. For result data, identifiers and locations are often indirectly linked, e.g. using DOIs that resolve to data sets, such as provided by Dryad and FigShare or domain-specific identifiers such as NCBI SRA project numbers.

#### 9. Provide enough metadata

In the course of a research project, data changes hands repeatedly and any additional information about it that is not explicitly recorded somewhere may be lost in the process. **Metadata, i.e. *data about data* that captures the who, what, where, when, why and how of a data set, may start to come into existence long before a sequencing experiment takes place yet may be needed long after the experiment has concluded and the results have been deposited in long-term repositories. Careful recording of metadata may prevent many unforeseen problems along this lifecycle**. Recent years have seen the development of checklists for the minimal amount of information required to describe a sequencing experiment, resulting in the guidelines MIxS [20], used in submissions to the European Nucleotide Archive and the European Genome-Phenome Archive, and MINSEQE [21], used in submissions to the INSDC’s Short Read Archive and ArrayExpress. However, these guidelines specify so few details that compliance to them by themselves does not suffice.

Among the metadata that may be required might be information for validating the project data *per se*, e.g. by identifying the data locations, their data formats (and dialects!), the provenance trail that resulted in the data - such the logging output of analysis pipelines-and file checksums. One step up from that, metadata should also capture the parameters of the assay (e.g. lab protocols, insert sizes, read lengths, primers, adaptors) to be able to perform meaningful analysis on the data. Then, across assays, the metadata about the study (e.g. the study subjects, their source(s), sampling methodologies, any treatments or manipulations) and the overarching investigative project (e.g. contact details of persons and labs involved, their access rights) should be recorded. The Investigation-Study-Assay tab-separated format, ISA-TAB [22], provides a convenient framework for capturing all of this information in spreadsheets that require no bioinformatics expertise yet are flexible enough to be able to incorporate the semantics, i.e. the explicit definition of meaning, of a lot of metadata by way of formal ontologies, whose adoption is growing rapidly [23].

### Future Proofing

#### 10. Be agile

High-throughput sequencing technologies (chemistry, sensors) are rapidly evolving, and analyses are becoming more complex, moving the field from essentially exploratory to hypothesis testing on multimodal data. In the process, multiple sequencing platforms are routinely combined (e.g. long reads for scaffolding with short reads for error correction). In light of these developments, the pragmatic approach is to be agile: although sequencer vendors provide their own data management and analysis platforms, **avoid vendor lock-in in proprietary file formats, databases, analysis tools, data access patterns or LIMSs**.

The advent of lightweight virtualization and “container” technologies give the flexibility to bring a multitude of complex analysis workflows to data libraries, provided common data standards are adhered to and the data infrastructure does not unduly limit the ways in which data can be accessed. “Containerized” deployment of analysis workflows on distributed compute nodes may allow large analyses to be parallelized more easily.

## Conclusions

Industry outlooks for the next five years suggest that the relative cost of HTS data management and data reduction will drop, but that the relative cost of downstream analysis will rise (Figure 2). To realize the relatively lower cost of data management and data reduction, sensible policies need to be established in order to cope effectively with the fact that HTS data generation outpaces innovation in data storage. This document attempts to establish some simple rules to guide the development of these policies at the levels of master data management, data analysis, and data sharing. Hard and fast rules are impossible to establish given the heterogeneity in technologies and institutional infrastructure and requirements, but this document does suggest some aspects of a generalized scenario, which is visualized as a flowchart in Figure 5.

**Figure 5.**
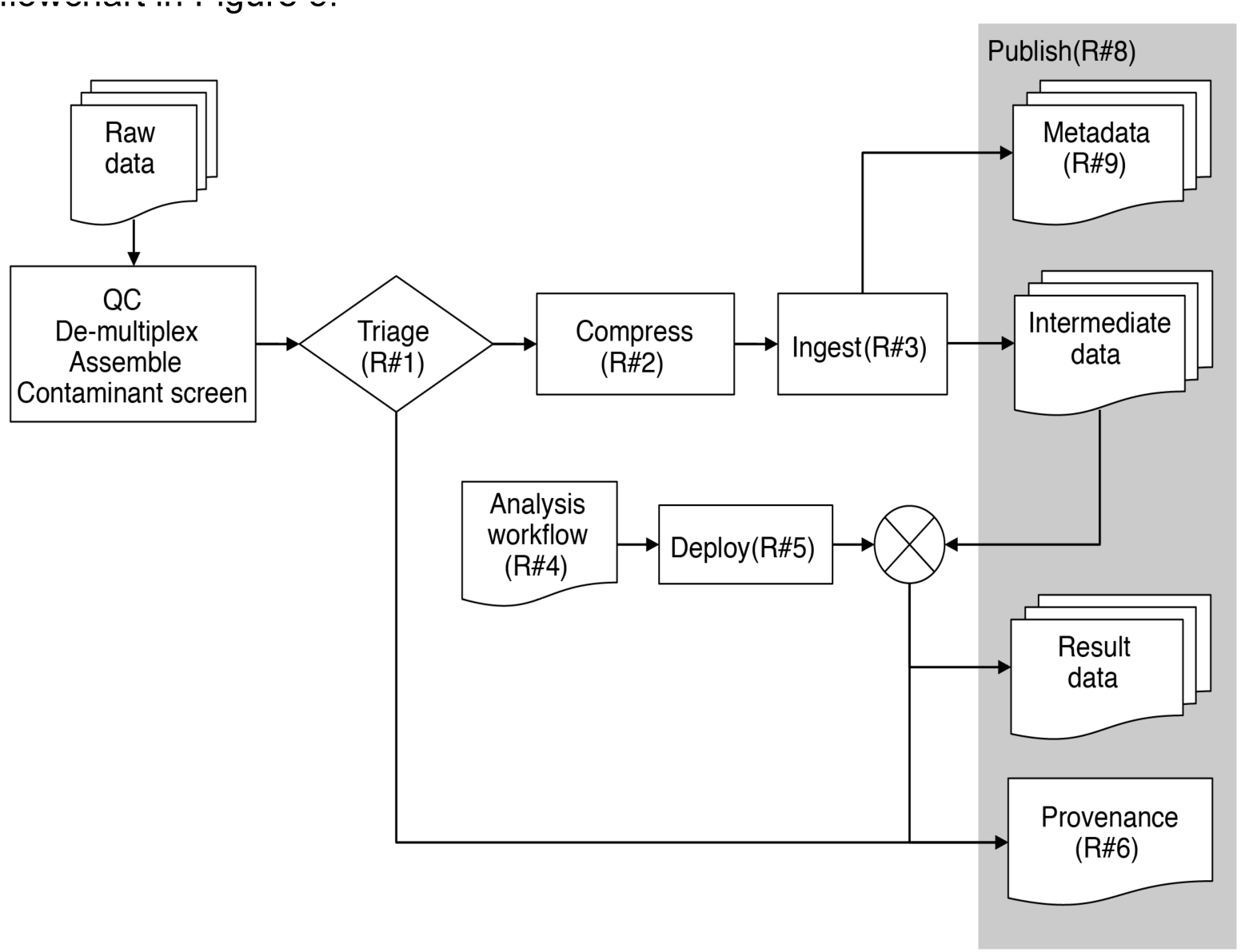
*Scenario for managing HTS data. Numbers between parenthese refer to discussed rules*.

In the scenario depicted in Figure 5, data undergo a planned series of movements and processes resulting in well-documented outcomes that can be shared among collaborators. The first steps in this process concern the generation of quality controlled, reduced and compressed intermediate data (e.g. BAM files) whose provenance and metadata are made available with the data through a publishing platform in which all persistently identifiable and locatable.

An arbitrary number of subsequent analysis steps is performed by processing these intermediate data with automated analysis workflows, ideally minimizing the movement and copying of the intermediate data (i.e. bringing the analysis to the data rather than vice versa). Each of these processes generates result data which again are ingested into a publishing platform. In the process, a paper trail of provenance is generated that extends the trail initiated during the earliest stages of data processing.

The scenario given here is necessarily very generic because the HTS field is evolving very rapidly, and specific use cases differ greatly among (and within) “accidental sequencing centers”. As such, perhaps the most important recommendation that can be made is to be agile in developing workflows, to think in terms of loosely-coupled modules that can be swapped in and out as needed, by avoiding proprietary platforms and technologies and pragmatically choosing interoperable standards instead.

## Footnotes

1 Data Management International: http://www.dama.org/i4a/pages/index.cfm?pageid=3339

2 e.g. see http://blogs.princeton.edu/onpopdata/2012/03/12/data-life-cycle/

3 http://dans.knaw.nl

4 http://datadryad.org

5 http://www.insdc.org/

6 e.g. see http://www.dcc.ac.uk/sites/default/files/documents/publications/DCCLifecycle.pdf

7 http://www.slideshare.net/gcoates/sharing-data-sanger-experiences

8 http://www.slideshare.net/gcoates/nextgeneration-sequencing-data-mangement

9 http://www.genengnews.com/gen-articles/managing-data-from-next-gen-sequencing/2449/

10 http://samtools.github.io/htsjdk/

11 http://www.ncbi.nlm.nih.gov/books/NBK47537/#File_Format_Guide_B.Overview_of_Input_Fo

12 http://www.ncbi.nlm.nih.gov/Traces/trace.cgi?cmd=show&f=formats&m=doc&s=format#sff

13 http://irods.org/

14 https://usegalaxy.org/

15 https://aws.amazon.com/s3/

16 http://ceph.com/ceph-storage/object-storage/

17 http://www.vagrantup.com/

18 https://puppetlabs.com/

19 https://www.docker.com/

20 https://virtualenv.pypa.io

21 For example: Galaxy (http://galaxyproject.org/), Taverna (http://www.taverna.org.uk/), eHive (http://www.ensembl.org/info/docs/eHive/index.html)

22 For example, make-like tools such as SnakeMake (https://bitbucket.org/johanneskoester/snakemake/wiki/Home)

23 For example the statistics language R (http://www.r-project.org/)

24 For example, Python (https://www.python.org/), Ruby (https://www.ruby-lang.org/en/) or Perl (http://www.perl.org/)

25 https://www.docker.com/

26 https://virtualenv.pypa.io/en/latest/

27 http://www.git-scm.com/

28 https://subversion.apache.org/

29 e.g. http://bitbucket.org, http://github.com, http://sourceforge.net

30 http://galaxyproject.org

31 for a good example of this, see https://usegalaxy.org/u/aun1/p/windshield-splatter

32 http://git-annex.branchable.com/

33 http://git-annex.branchable.com/special_remotes/

34 http://www.slideshare.net/gcoates/nextgeneration-sequencing-data-mangement

35 http://www.slideshare.net/gcoates/nextgeneration-sequencing-data-mangement

## References

1. “Inter-university Consortium for Political and Social Research (ICPSR)” (2012) Guide to Social Science Data Preparation and Archiving: Best Practice Throughout the Data Life Cycle. 5th ed. Ann Arbor, MI.

2. Lawrence B, Jones C, Matthews B, Pepler S, Callaghan S (2011) Citation and Peer Review of Data: Moving Towards Formal Data Publication. Int J Digit Curation 6: 4–37. doi:10.2218/ijdc.v6i2.205.

3. Clark T, Ciccarese PN, Goble CA (2013) Micropublications: a Semantic Model for Claims, Evidence, Arguments and Annotations in Biomedical Communications.

4. Vlieg J de, van Schaik R, Aerts P, Lusher S, Sienstra F, et al. (2013) Data-Stewardship in the Big Data Era: Taking Care of Data. Amsterdam. 9 p.

5. Sboner A, Mu XJ, Greenbaum D, Auerbach RK, Gerstein MB (2011) The real cost of sequencing: higher than you think! Genome Biol 12: 125. doi:10.1186/gb-2011-12-8-125.

6. Cock PJA, Fields CJ, Goto N, Heuer ML, Rice PM (2010) The Sanger FASTQ file format for sequences with quality scores, and the Solexa/Illumina FASTQ variants. Nucleic Acids Res 38: 1767–1771.

7. Wan R, Anh VN, Asai K (2012) Transformations for the compression of FASTQ quality scores of next-generation sequencing data. Bioinformatics 28: 628–635. doi:10.1093/bioinformatics/btr689.

8. Bonfield JK, Mahoney M V (2013) Compression of FASTQ and SAM format sequencing data. PLoS One 8: e59190.

9. Deorowicz S, Grabowski S (2011) Compression of DNA sequence reads in FASTQ format. Bioinformatics 27: 860–862.

10. Li H, Handsaker B, Wysoker A, Fennell T, Ruan J, et al. (2009) The Sequence Alignment/Map format and SAMtools. Bioinformatics 25: 2078–2079. doi:10.1093/bioinformatics/btp352.

11. Hsi-Yang Fritz M, Leinonen R, Cochrane G, Birney E (2011) Efficient storage of high throughput DNA sequencing data using reference-based compression. Genome Res 21: 734–740. doi:10.1101/gr.114819.110.

12. Loh P-R, Baym M, Berger B (2012) Compressive genomics. Nat Biotechnol 30: 627–630. doi:10.1038/nbt.2241.

13. Jordan C, Stanzione D, Ware D, Lu J, Noutsos C (2010) Comprehensive data infrastructure for plant bioinformatics. 2010 IEEE Int Conf Clust Comput Work Posters (CLUSTER Work: 1–5. doi:10.1109/CLUSTERWKSP.2010.5613093.

14. Lampa S, Dahlö M, Olason PI, Hagberg J, Spjuth O (2013) Lessons learned from implementing a national infrastructure in Sweden for storage and analysis of next-generation sequencing data. Gigascience 2: 9. doi:10.1186/2047-217X-2-9.

15. Chiang G-T, Clapham P, Qi G, Sale K, Coates G (2011) Implementing a genomic data management system using iRODS in the Wellcome Trust Sanger Institute. BMC Bioinformatics 12: 361.

16. Noble WS (2009) A Quick Guide to Organizing Computational Biology Projects. PLoS Comput Biol 5. doi:10.1371/journal.pcbi.1000424.

17. Sandve GK, Nekrutenko A, Taylor J, Hovig E (2013) Ten Simple Rules for Reproducible Computational Research. PLoS Comput Biol 9: e1003285.

18. Szalay A, Gray J (2006) 2020 computing: science in an exponential world. Nature 440: 413–414. doi:10.1038/440413a.

19. Stein LD (2010) The case for cloud computing in genome informatics. Genome Biol 11: 207.

20. Yilmaz P, Kottmann R, Field D, Knight R, Cole JR, et al. (2011) Minimum information about a marker gene sequence (MIMARKS) and minimum information about any (x) sequence (MIxS) specifications. Nat Biotechnol 29: 415–420. doi:10.1038/nbt.1823.

21. MINSEQE: Minimum Information about a high-throughput Nucleotide SeQuencing Experiment - a proposal for standards in functional genomic data reporting (2012). 0. Available: http://www.fged.org/site_media/pdf/MINSEQE_1.0.pdf.

22. Rocca-Serra P, Brandizi M, Maguire E, Sklyar N, Taylor C, et al. (2010) ISA software suite: supporting standards-compliant experimental annotation and enabling curation at the community level. Bioinformatics 26: 2354–2356. doi:10.1093/bioinformatics/btq415.

23. Katayama T, Wilkinson MD, Aoki-Kinoshita KF, Kawashima S, Yamamoto Y, et al. (2014) BioHackathon series in 2011 and 2012: penetration of ontology and linked data in life science domains. J Biomed Semantics 5: 5. doi:10.1186/2041-1480-5-5.

